# Maternal immune activation accelerates puberty initiation and alters mechanical allodynia in male and female C57BL6/J mice

**DOI:** 10.1101/2022.01.13.476235

**Authors:** Xin Zhao, Mary Erickson, Ruqayah Mohammed, Amanda C. Kentner

## Abstract

The mechanisms that link maternal immune activation (MIA) with the onset of neurodevelopmental disorders remain largely unclear. Accelerated puberty is also associated with a heightened risk for psychopathology in later life but there is a dearth of evidence on the impacts of maternal infection on pubertal timing. We examined the effects of MIA on reproductive development, mechanical allodynia, and sensorimotor gating in juvenile, adolescent, and adult male and female mice. Moreover, we investigated hypothalamic neural markers associated with the reproductive and stress axes. Finally, we tested the mitigating effects of environmental enrichment (EE), which has clinical relevancy in human rehabilitation settings. Our results show that administration of polyinosinic-polycytidylic acid (poly(I:C)) on gestational day 12.5 led to early preputial separation, vaginal openings, and age of first estrus in offspring. MIA exposure altered pain sensitivity across development and modestly altered prepulse inhibition. The downregulation of *Nr3c1* and *Oprk* mRNA in the hypothalamus of juvenile mice suggests that MIA’s effects may be mediated through disruption of hypothalamic-pituitary-adrenal axis activity. In contrast, life-long housing with EE rescued many of these MIA-induced consequences. Overall, our findings suggest that accelerated puberty may be associated with the deleterious effects of infection during pregnancy and the onset of psychopathology.

## 1. Introduction

Puberty is an important developmental stage marked by dramatic biological transitions. Altered pubertal timing (i.e., early puberty onset) has been linked with a heightened risk for psychopathology and behavioral disruptions later in life (Colich et al., 2020; Dimler and Natsuaki, 2015; Hamlat et al., 2019), and recent findings have shown a higher incidence rate of precocious puberty in people diagnosed with autism (Corbett et al., 2020; Geier and Geier, 2021). Moreover, a growing number of studies in humans suggest that the acceleration of various physiological metrics of development are associated with early life adversities (ELA; Belsky et al., 2015; Berghänel et al., 2017; Colich et al., 2020; Gur et al., 2019; Miller et al., 2020; Sumner et al., 2019). In rodent studies, maternal and neonatal immune challenge, in addition to maternal separation have each accelerated puberty development (Cakan et al., 2018; Grassi-Oliveira et al., 2016; Yano et al., 2020). Given the abovementioned associations, accelerated pubertal timing may play an important part in mediating the effects of ELA on physical and mental health problems such as chronic pain and disrupted sensorimotor gating which tend to occur as an individual matures and develops in an age-specific manner (El Tumi et al., 2017; Romero et al., 2010).

Exposure to a viral or bacterial infection in utero is an ELA. A wealth of epidemiological evidence has revealed a link with these infections, known as maternal immune activation (MIA), and the later life emergence of psychiatric disorders (for review see Estes and McAllister, 2016) which are associated with disrupted mechanical allodynia/pain or tactile sensory processing (Schaffler et al., 2019). However, it remains largely unclear what impact MIA may have on pubertal timing. Preclinical rodent MIA models have been widely used to mimic infection in humans in order to investigate their underlying pathogenic mechanisms and potential therapeutic interventions (Estes and McAllister, 2016; Kentner et al., 2019a; Knuesel et al., 2014). One of the most commonly used immunogens in rodent MIA models is polyinosinic:polycytidylic acid (poly (I:C)), a commercially available synthetic analog of double-stranded RNA that has been shown to induce an extensive collection of innate immune responses (Mueller et al., 2019). These responses lead to a wide array of abnormalities in brain morphology (Li et al., 2009; Meyer et al., 2006), in addition to neurochemical, and pharmacological reactions (Zuckerman et al., 2003; Zuckerman and Weiner, 2005) that are associated with altered behaviors and cognitive abilities (Li et al., 2009; Ozawa et al., 2006; Zhao et al., 2021a; Zhao et al., 2021b). Using the poly (I:C) model, one recent study in rats demonstrated that immune challenge at gestational day 12, but not 14, advanced the appearance of vaginal opening in females (Cakan et al., 2018). However, it remains unclear the underlying mechanisms and whether gestational immune challenge affects pubertal timing in males.

Normal puberty development requires the integration of multiple molecular and cellular signals along the hypothalamus-pituitary-gonadal axis. This includes the stimulatory effects of kisspeptin signaling on gonadotropin-releasing hormone (GnRH) release (Han et al., 2005; Herbison et al., 2010) and neurokinin B signaling (including the receptor encoded by *Tac3r*; Navarro, 2020; Navarro et al., 2012; Topaloglu et al., 2009) which initiates a series of endocrine changes that drives the development of sex characteristics (Parent et al., 2003). In parallel, the expression level of the GnRH and kisspeptin genes increase across the initiation of puberty (Gore et al., 1996; Han et al., 2005; Parfitt et al., 1999). Previous evidence has also shown that disruptions in the gene Makorin ring finger 3 (*Mkrn3*) are associated with premature sexual development (Abreu et al., 2013; Abreu et al., 2015). MIA may therefore alter sexual maturation by affecting the expression of genes associated with these signals in the hypothalamus.

According to the life history theory, early life adversities represent a deviation from the expected environment (Belsky, 2019) and the accelerated pubertal timing may therefore maximize reproductive fitness prior to mortality. This theory highlights a critical role of the environment in shaping phenotypic development. It also implies that manipulating the environment in which an individual lives could be an interventional strategy. In contrast to an adverse early environment, opportunities for enhanced putatively positive experiences in early life, such as living in an enriched environment (EE) has been shown to confer a number of beneficial effects including enhanced brain plasticity (e.g., increased dendritic branching, synaptogenesis, neurogenesis; (Brenes et al., 2016; Kolb et al., 1998; Van Praag et al., 2000) as well as improvement in cognitive functions (Williams et al., 2001; Zeleznikow-Johnston et al., 2017). Our lab has previously demonstrated that EE rescues disruptions to social motivation, spatial discrimination, and neural markers associated with synaptic plasticity and stress responses following MIA in both mice and rats (Connors et al., 2014; Kentner et al., 2016; Núñez Estevez et al., 2020; Zhao et al., 2021a; Zhao et al., 2020; Zhao et al., 2021b). These findings underscore EE’s potential in the mitigation of the behavioral and neurophysiological impairments associated with ELA.

In the current study, pubertal timing (vaginal opening in females and complete preputial separation in males) and estrous cyclicity of mouse offspring exposed to either prenatal poly (I:C) or saline were evaluated. We also examined mechanical allodynia and sensorimotor gating at different developmental ages (juvenile, adolescence, and adulthood) to evaluate whether these physiological functions were altered as a function of reproductive status. Furthermore, we evaluated male and female offspring since the two sexes tend to respond differently in terms of thresholds to painful stimuli and sensorimotor gating (Nicotra et al., 2014; Swerdlow et al., 1999). In addition to these behavioral measurements, we also quantified mRNA expression of genes associated with the endogenous opioid system, gamma-aminobutyric acid (GABA)ergic signaling and stress responses, which have been implicated in modulating mechanical allodynia and sensorimotor gating (Deslauriers et al., 2013; Ossipov et al., 2010; Yeomans et al., 2010). We selected opioid receptor, kappa 1 (*Oprk)* gene because a previous clinical study demonstrated that polymorphisms in *Oprk* but not mu- and delta-opioid receptor genes is associated with experimental pain sensitivity (Sato et al., 2013). We anticipate that our findings will shed light on the mechanisms underlying the deleterious effects of gestational immune insults and will give rise to new insights into possible intervention strategies following ELA.

## 2. Materials and Methods

### 2.1. Animals and Housing

Male and female C57BL/6J mice from the Jackson Laboratory (Bar Harbor, ME) were housed on a 12 h light/dark cycle (0700–1900 light) maintained at 20°C. Animals had full access to food and water. **Figure 1** outlines the experimental procedures used in this study. Animals were pair-housed in one of two types of cages: environmental enrichment (EE; N40HT mouse cage, Ancare, Bellmore, NY), comprised of a larger cage and access to toys, tubes, a Nylabone, Nestlets® (Ancare, Bellmore, NY) and Bed-r’Nest® (ScottPharma Solutions, Marlborough MA), or standard cages (SD; N10HT mouse cage, Ancare, Bellmore, NY) with Nestlets® and a Nylabone only. Male animals were paired in SD conditions unless they were breeding (1 male: 2 female design), during which they could be in an EE cage. Female mice were individually housed in a fresh clean cage immediately after breeding, maintaining their assigned housing condition.

**Figure 1.**
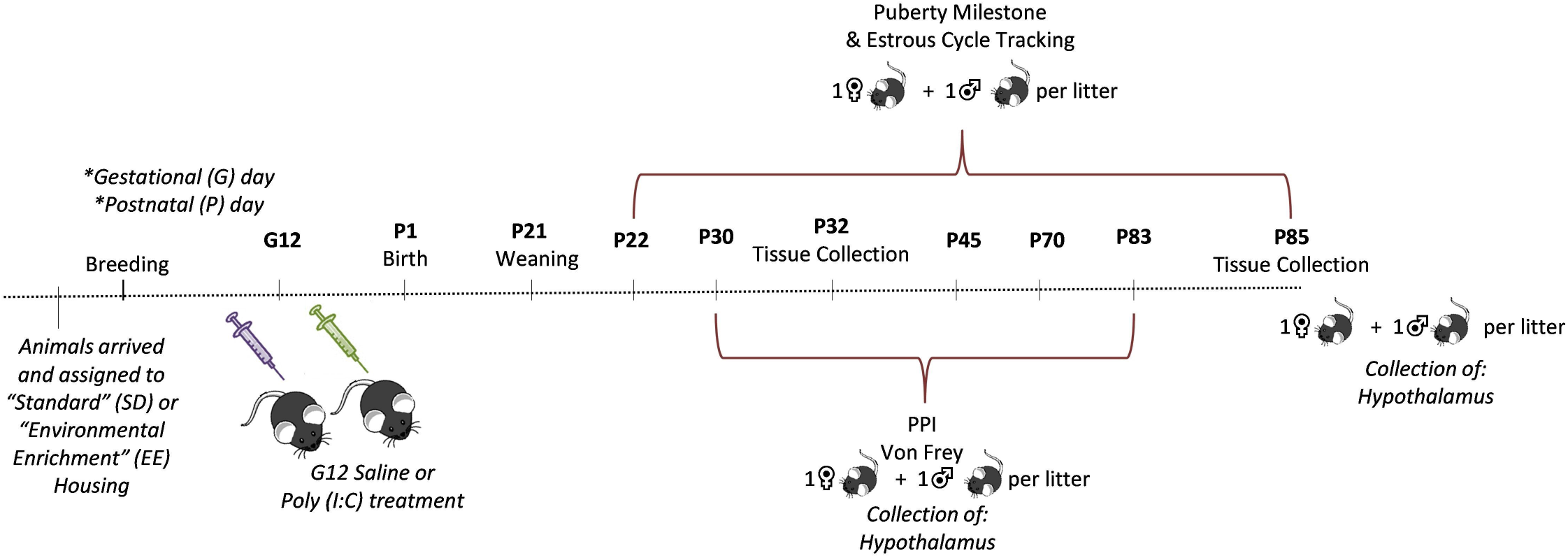
Experimental Timeline.

Pregnant dams were randomly allocated to receive either 20 mg/kg intraperitoneal injection of polyinosine-polycytidylic acid (poly (I:C); tlrl-picw, lot number PIW-41-03, InvivoGen), or vehicle (sterile pyrogen-free 0.9% NaCl) on the morning of gestational day (G) 12. Due to the considerable occurrence of spontaneous abortion induced by high molecular weight poly (I:C) (Mueller et al., 2019), we choose low molecular weight poly(I:C). The innate immune response of this dosage was verified in other studies (McGarry et al., 2021; Mueller et al., 2019), and we have also demonstrated that 20 mg/kg of low molecular weight poly (I:C) induced robust body weight loss (Zhao et al., 2021a). Toys, tubes, and bones were removed from cages on G19, to ensure pup safety, and returned on postnatal day (P)15. Additional details can be found in the methodological reporting table from Kentner et al. (2019a), provided as **Supplementary Table S1**. The Institutional Animal Care and Use Committee at the Massachusetts College of Pharmacy and Health Sciences approved all animal procedures which were carried out in compliance with the recommendations outlined by the Guide for the Care and Use of Laboratory Animals of the National Institutes of Health.

### 2.3. Offspring Behavior

#### 2.3.1. Experimental Design

On P21, offspring were weaned into same-sex groups, keeping their original housing assignments (SD = 2-3 animals/cage; EE = 4-5 animals/cage). No more than one male and one female animal from each litter were evaluated on each endpoint. Male and female offspring from one set of animals (juvenile time point) were evaluated for prepulse inhibition (PPI) and then the von Frey test on P30. Tissue collection took place on P32. Separate sets of animals were evaluated on the timing of puberty milestones, PPI, and then the von Frey test during adolescence and adulthood A second tissue collection time point took place on P85.

#### 2.3.2. Puberty Markers and Estrous Cycle

Starting on the day of weaning (P21), a subset of male and female mice (n = 9-13) was evaluated for age of complete balano-preputial separation and vaginal opening (closed or complete), respectively (Grassi-Oliveira et al., 2016). Vaginal swabs were collected daily to track the estrous cycle. Estrous cycle data was evaluated to determine each animal’s age when they were in the estrus phase of their cycle for the first time, their total number of full estrous cycles, and the average length of their estrous phase across the study. No behavioral endpoints were assessed until after puberty in these animals.

#### 2.3.3. Prepulse Inhibition of the Acoustic Startle Reflex

On P45, P70, and P83, prepulse inhibition (PPI) of the acoustic startle reflex was evaluated (n = 8-13) using six startle chambers from San Diego Instruments (San Diego Instruments, San Diego, CA, USA). Boxes and equipment were cleansed with the disinfectant Quatriside-TB (Parmacal Research Laboratories, Inc) and dried between each animal and session. Each session followed the protocol described by Giovanoli et al. (2013). In brief, the trial consisted of pulse-alone, prepulse-plus-pulse and prepulse-alone trials, as well as no-stimulus trials in which only a background noise of 65 dB was presented. In the pulse-alone trial, a 40-ms pulse of white noise (120 dB) was administered. In the prepulse-plus-pulse trial, a 20-ms burst of white noise at five different intensities (69, 73, 77, 81, 85 dB) corresponding to 4, 8, 12, 16, and 20 dB above the background noise was administered prior to the 120 dB white noise. The interval between the prepulse and pulse during the prepulse-plus-pulse trials was 100 ms. Only a 20-ms burst of the 5 different prepulses was administered during the prepulse-alone trials. Each session was initiated by 6 pulse-alone trials. Then, each prepulse-plus-pulse, prepulse-alone, pulse-alone and no-stimulus trial was presented 12 times in a pseudorandom order. The session terminated with 6 consecutive pulse-alone trials. The average interval between successive trials (ITI) was 15 ± 5 sec. For each animal and at each of the five possible prepulse intensities, PPI was calculated as percent inhibition of startle response obtained in the prepulse-plus-trials compared to pulse-alone trials: [1-(mean reactivity on prepulse-plus-pulse trials / mean reactivity on pulse-alone trials) × 1/100] and then expressed as %PPI.

#### 2.3.4. Mechanical Allodynia

Evaluation of mechanical allodynia commenced 3 hours following completion of PPI testing on P30, P45, or P70 (n = 7-13). Each animal was placed inside an acrylic cage, with a wire grid floor. This acclimatization period took place for 30 minutes prior to the test session. Mechanical allodynia was evaluated with a pressure-meter which consisted of a hand-held force transducer fitted with a polypropylene rigid tip (Electronic von Frey Aesthesiometer, IITC, Inc, Life Science Instruments, Woodland Hills, CA, USA). The polypropylene tip was applied perpendicularly to the central area of the left hind paw with increasing pressure. The trial terminated when the animal withdrew their paw from the tip. Stimulus intensity was recorded by the electronic pressure-meter when the paw was withdrawn. The average of four test trials was calculated as the mechanical withdrawal threshold (grams) for each session (Yan and Kentner, 2017).

### 2.4. Tissue collection

On the morning of either P32 or P85 (n = 6-8), a mixture of ketamine/xylazine (150 mg/kg, i.p/15 mg/kg, i.p) was used to anesthetize animals. Animals were perfused intracardially with a chilled phosphate buffer solution. Hypothalamus was dissected, frozen on dry ice, and stored at −75°C until processing.

### 2.5. RT-PCR

The RNeasy Lipid Tissue Mini Kit (Qiagen, 74804) was used to extract RNA from frozen tissue. The extraction was then eluted in RNase-free water. A NanoDrop 2000 spectrophotometer (ThermoFisher Scientific) was used to quantify the isolated RNA. Total RNA was reverse transcribed to cDNA using the Transcriptor High Fidelity cDNA Synthesis Kit (#5081963001, Millipore Sigma) and stored at -20°C until analysis. Quantitative real-time PCR with Taqman™ Fast Advanced Master Mix (#4444963, Applied Biosystems) was used to measure the mRNA expression of kisspeptin (*Kiss1*, Mm03058560_m1), *Kiss1* receptor (*Kiss1r*, Mm00475046_m1), glucocorticoid receptor (*Nr3c1*, Mm00433832_m1), gonadotropin-releasing hormone 1 (*Gnrh1*, Mm01315605_m1), tachykinin 2 (*Tac2*, Mm01160362_m1), tachykinin receptor 3 (*Tac3r*, Mm00445346_m1), gamma-aminobutyric acid (GABA) A receptor, subunit gamma 2 (*Gabrg2*, Mm00433489_m1), makorin, ring finger protein, 3 (*Mkrn3*, Mm00844003_s1), opioid receptor, kappa 1 (*Oprk*, Mm01230885_m1), and FK506 binding protein 5 (*Fkbp5*, Mm00487401_m1). Samples were run in triplicate using optical 96-well plates (Applied Biosystems StepOnePlus™ Real-Time PCR System) and relative gene expression levels were evaluated using the 2 –_ΔΔ_^CT^ method (Livak and Schmittgen, 2001). Gene expression was normalized to 18S (Hs99999901_s1) mRNA levels, which have been used as endogenous control for gene expression assay in previous MIA studies (Meyer et al., 2008; Smith et al., 2020). Data are presented as mean expression relative to same sex SD-saline treated controls.

### 2.6. Statistical analysis

Statistics were performed using the Statistical Software for the Social Sciences (SPSS) version 26.0 (IBM, Armonk, NY) or GraphPad Prism (version 9.0). Time to vaginal opening, balano-preputial separation, and age of first estrus stage were assessed using the Log-rank (Mantel Cox) test and plotted as a survival curve. Following the overall omnibus test for each measure, individual Log-rank tests were conducted as post hocs, alongside a Bonferonni alpha correction to protect against multiple comparisons. The Shapiro-Wilk test was used to assess the assumption of normality and Kruskal-Wallis tests (expressed as *X*^2^) employed in rare cases of significantly skewed data. Since different groups of animals were used across the evaluation time points, three-way ANOVAs (Sex x MIA x Housing) were used in the evaluation of PPI and mechanical allodynia, with body weight included as a covariate for the latter measure. MIA x Housing interactions and main effects were further analyzed in each sex separately (Gildawie et al., 2021), ensuring that each analysis was performed with only one animal from each litter. Since the qPCR data were normalized to same sex controls, a two-way ANOVA (MIA x Housing) was conducted for each gene of interest in each sex separately. Tukey’s post hocs were applied except where there were fewer than three levels, in which case pairwise t-tests and Levene’s (applied in the occurrence of unequal variances) were utilized. Given that different sets of animals were evaluated on the major endpoints of this study, Pearson correlations were only analyzed between age of vaginal opening, first estrus, preputial opening, and mechanical allodynia (P45, P70), PPI (P45, P70, P83), and hypothalamic gene expression (P85). The False Discovery Rate (FDR) was applied to correct for multiple testing procedures in all gene expression experiments. Data are graphically expressed as mean ± SEM. The partial eta-squared (*n*^*p*2^) is also reported as an index of effect size for the ANOVAs (the range of values being 0.02 = small effect, 0.13 = moderate effect, 0.26 = large effect; Miles and Shevlin, 2001).

## 3. Results

### 3.1. MIA challenge accelerates puberty initiation

For female animals, there was a MIA by housing interaction for day of vaginal opening (F(1, 44) = 7.591, p = 0.009, *n*^*p*2^ = 0.187; **Figure 2A**). SD-poly (I:C) animals displayed vaginal openings significantly earlier than SD-saline mice (p = 0.010). There was no significant difference between EE-saline and EE-poly (I:C) female mice (p>0.05), suggesting that EE housing was modestly protective against the accelerated puberty. There was a significant effect on vaginal opening timing across all groups (*X2*(3) = 8.949, p = 0.03; **Figure 2B)**. Vaginal opening in SD-poly (I:C) mice occurred earlier compared to SD-saline (*X2* (1) = 11.53, p = 0.0007) and lack of statistical difference between EE-saline and EE-poly (I:C) animals (p>0.05).

**Figure 2.**
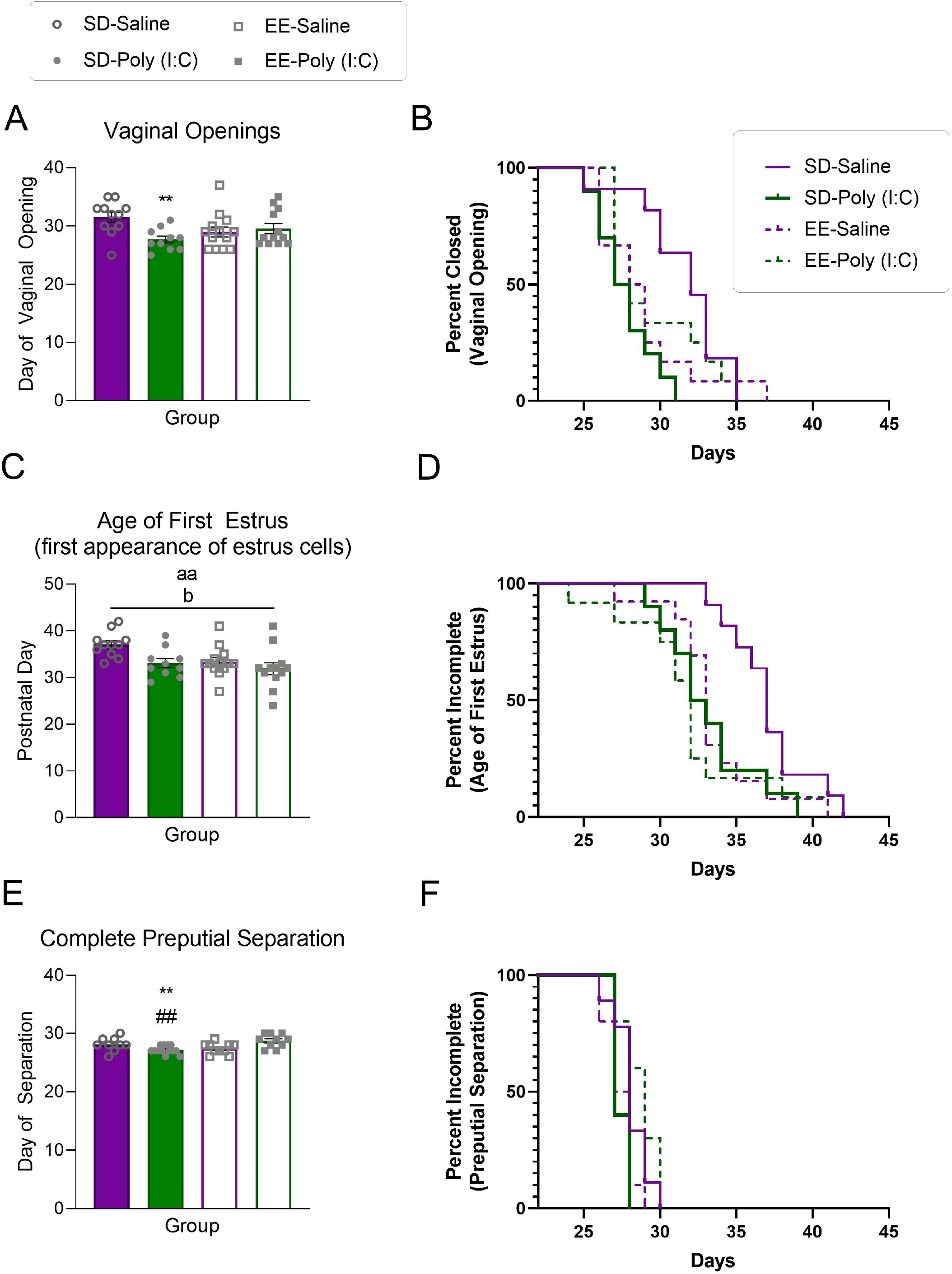
The effects of maternal immune activation (MIA) and environmental enrichment on reproductive milestones. **(A)** Day of vaginal opening, **(B)** survival curves of full vaginal opening for each group expressed across time, **(C)** day of first estrus, **(D)** survival curves of day of first estrus for each group expressed across time, **(E)** day of complete preputial separation, and **(F)** survival curves of complete balano-preputial separation for each group expressed across time. Data are expressed as mean ± SEM. *p < 0.05, **p < 0.01, ***p <0.001, versus SD-saline; #p < 0.05, ##p < 0.01, ###p <0.001, versus EE-poly (I:C); ap < 0.05, aap < 0.01, main effect of MIA; bp < 0.05, main effect of housing.

A main effect of MIA was found for age of first estrus (F(1, 44) = 7.188, p = 0.010, *n*^*p* 2^ = 0.146; **Figure 2C**) in that poly (I:C) females reached this stage earlier than saline treated controls (p = 0.019). A main effect of housing (F(1, 44) = 5.767, p = 0.021, *n*^*p* 2^ = 0.121) revealed that EE females displayed their first estrus sooner than SD housed mice (p = 0.026). The overall Log-rank test was significant across all groups (*X2* (3) = 8.671, p = 0.034; **Figure 2D**). SD-poly (I:C) females had their first estrus sooner than SD-saline females ((*X2* (1) = 8.038, p = 0.005) and the same pattern was maintained between EE-saline and SD-saline females ((*X2* (1) = 6.535, p = 0.011). There were no differences across groups in the number of full estrous cycles nor in the average length of the estrous cycle across the study (p >0.05).

For male animals, a MIA by housing interaction was also revealed for day of complete preputial separation (F(1, 40) = 7.421, p = 0.010, *n*^*p* 2^ = 0.163; **Figure 2E)**. SD-poly (I:C) males displayed complete preputial separation earlier than SD-saline (p = 0.009) and EE-poly (I:C) mice (p = 0.001). The overall Log-rank test was significant across all groups (*X2* (3) = 10.99, p = 0.012; **Figure 2F**). SD-poly (I:C) males had complete preputial separation earlier than EE-poly ((I:C); *X2* (1) = 8.038, p = 0.005) but not SD-saline males ((*X2* (1) = 3.723, p = 0.054). EE-poly (I:C) males also took longer for complete preputial separation to occur compared to EE-saline males ((*X*2 (1) = 6.102, p = 0.014).

### 3.2. MIA challenge affects mechanical allodynia in an age dependent manner

There was no significant interaction (Sex x MIA x Housing) for mechanical allodynia, as measured by the von Frey test on P30 (p >0.05; **Figure 3AB**), but when separated by sex, our results revealed a significant interaction (MIA x Housing) for both males (F(1, 32) = 7.769, p = 0.009, n_p_2 = 0.195) and females (F(1, 28) = 10.461, p = 0.003, *n*^*p* 2^ = 0.272). Male and female SD-poly (I:C) offspring had lower thresholds than SD-saline (males: p = 0.010; females: p = 0.026) and EE-poly (I:C) P30 offspring (males: p = 0.011; females: p = 0.043). Female EE-saline animals had lower thresholds compared to SD-saline mice (p=0.008) at this age. On P45, there were significant interactions (**Figure 3AB**) between MIA and housing for males (F(1, 38 = 12.524, p = 0.001, n^p2^ = 0.248) and females (F(1, 41) = 4.163, p = 0.048, *n*^*p*2^ = 0.092). Compared to SD-saline animals, SD-poly (I:C) male (p = 0.010) and female (p = 0.026) mice had higher thresholds. Male SD-poly (I:C) mice had elevated thresholds compared to EE-poly (I:C) housed animals (p = 0.011). Female EE-saline animals had higher thresholds than SD-saline (p = 0.008). At P70, there were no MIA or housing effects for mechanical allodynia (p>0.05); there was only a significant main effect of sex with males having higher thresholds than females (F(1, 82) = 5.009, p = 0.028, *n*^*p* 2^ = 0.058; **Figure 3AB**).

**Figure 3.**
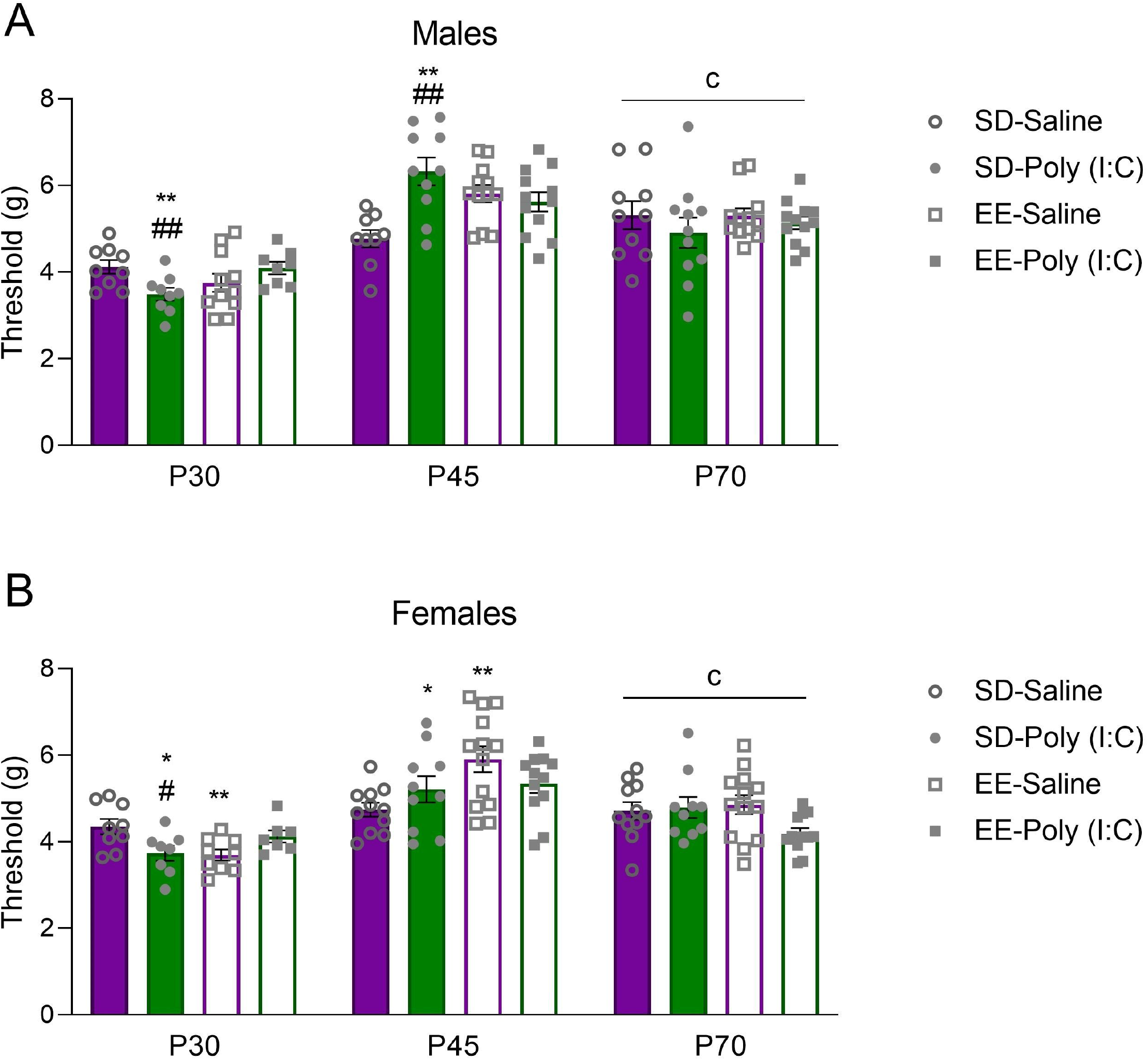
The effects of maternal immune activation (MIA) and environmental enrichment on mechanical allodynia. **(A)** Male and **(B)** Female mechanical paw withdrawal thresholds (grams) on the von Frey test. Data are expressed as mean ± SEM. Juvenile animals (P30): *p < 0.05, **p < 0.01, ***p <0.001, versus SD-saline; #p < 0.05, ##p < 0.01, ###p <0.001, versus EE-poly (I:C); cp < 0.05, main effect of sex.

### 3.3. Effect of MIA and environmental enrichment on prepulse inhibition (PPI)

Mice in all treatment groups, and across all ages, demonstrated increased % PPI as the intensity was raised from 69 to 85 dB. With respect to the P30 time point, there was no significant interaction (MIA x Housing x Intensity) identified (p>0.05; **Figure 4A**). However, PPI was disrupted as a function of MIA (males: F(1, 33) = 9.671, p = 0.004, *n*^*p* 2^ = 0.277; **Figure 4A**) and housing (females: F(1, 28) = 4.404, p = 0.045, *n*^*p*2^ = 0.136; **Figure 4A**) at the 69 dB intensity but follow-up tests were not significant (p>0.05). At the 73 dB intensity (male MIA x Housing: F(1, 33) = 8.495, p = 0.006, *n*^*p*2^ = 0.205), SD-poly ((I:C); p = 0.010) and EE-saline (p = 0.002) had higher PPI than SD-saline male mice. There were no sex, MIA, or housing effects at 77 dB (p> 0.05). At the 81 dB intensity, there was a three-way sex by MIA by housing interaction (F(1,62) = 5.623, p = 0.0251, *n*_p_ 2 = 0.083) where female EE-poly (I:C) mice had higher PPI values compared to EE-saline female mice (p = 0.038). Across all intensity values on P30, male EE mice had higher mean % PPI values than SD (p = 0.023; **Figure 4B**).

**Figure 4.**
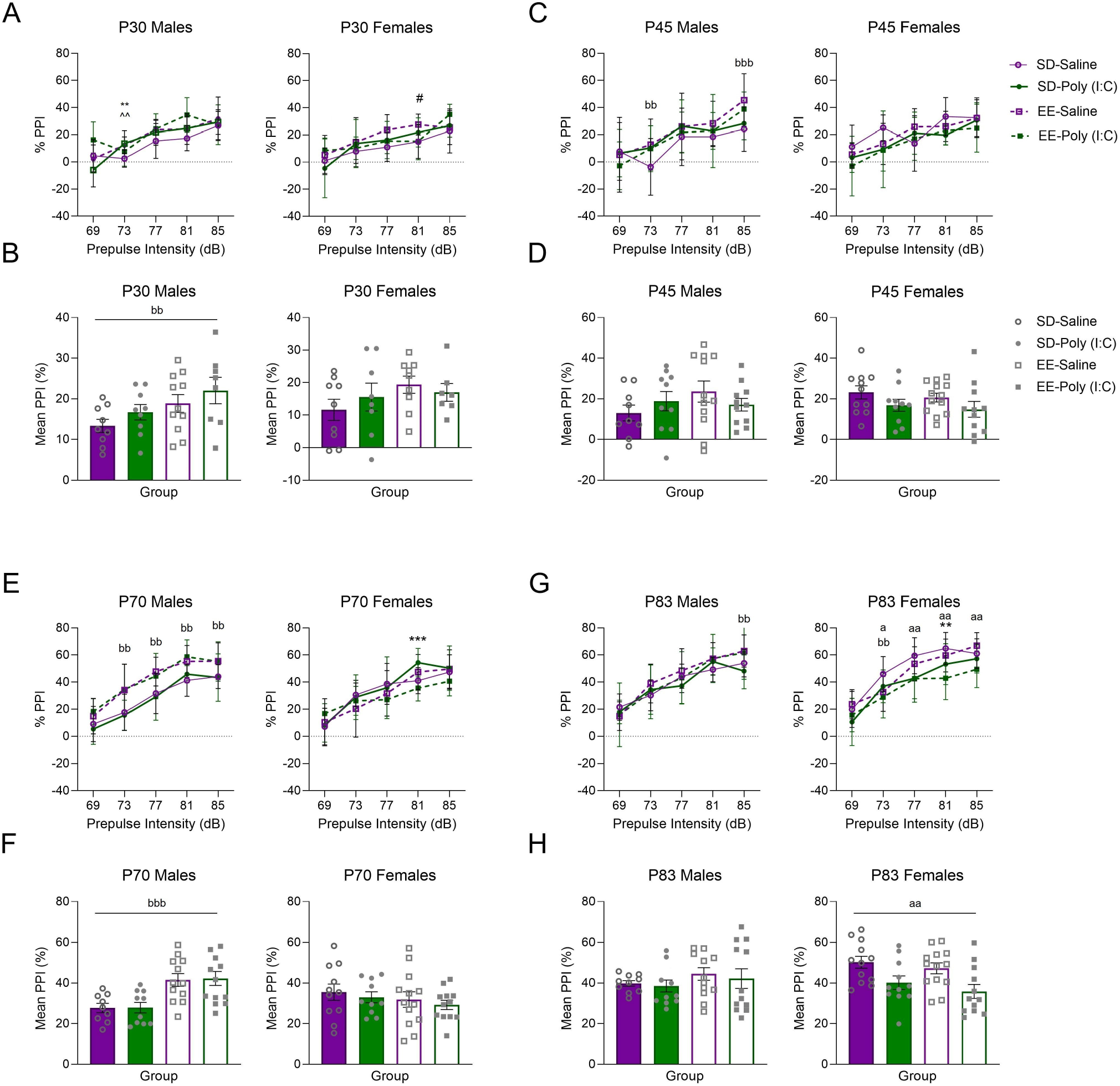
The effects of maternal immune activation (MIA) and environmental enrichment on prepulse inhibition of the acoustic startle reflex. The line plots display percent prepulse inhibition (PPI) as a function of increasing prepulse intensities for male (left) and female (right) mouse offspring on **(A)** postnatal day (P)30, **(C)** P45, **(E)** P70, and **(G)** P83. The bar plots show the mean percent prepulse inhibition across all five prepulse intensities for each of (B) P30, (D) P45, (F) P70, and (G) P83. Data are expressed as mean ± SEM. *p < 0.05, **p < 0.01, ***p <0.001, versus SD-saline vs SD-poly (I:C); #p < 0.05, EE-saline versus EE-poly (I:C); ^p < 0.05, SD-saline vs EE-saline; &p < 0.05, SD-poly (I:C) v EE-poly (I:C); aap < 0.01, aaa < 0.01, main effect of MIA; bbp < 0.01, bbbp < 0.01, main effect of housing.

At the P45 time point, there was no significant interaction (MIA x Housing x Intensity) identified (p >0.05; **Figure 4C**). However, there was a significant sex by housing interaction for the 73 dB (F(1, 81) = 6.913, p = 0.01, *n*^*p*2^ = 0.079; **Figure 4C**) and 85 dB intensities (F(1, 81) = 11.067, p = 0.001, *n*^*p*2^ = 0.120) where male EE animals had higher PPI than male SD (73 dB: p = 0.017; 85 dB: p = 0.0001). The mean % PPI across all intensity values revealed a sex by housing effect (F(1, 81) = 4.056, p = 0.047, *n*^*p* 2^ = 0.048**; Figure 4D**).

On P70, while the interaction (MIA x Housing x Intensity) was not significant (p >0.05; **Figure 4E**), there were sex by housing interactions across the individual 73-, 77-, 81-, and 85-dB intensities (p = 0.0001; **Figure 4E**). The mean % PPI across all intensity values confirmed this interaction (males: F(1, 41) = 24.563, p = 0.001, *n*^*p* 2^ = 0.375; females: N.S **Figure 4F**). At this age, male EE mice had a higher mean % PPI than SD male mice across all intensities. On P83, this sex by housing interaction was lost across all intensities combined (mean PPI (%); p >0.05; **Figure 4GH**) although sex by housing effects were still observed at the individual 73 and 85, dB intensities (males 85 dB: p = 0.005; females 73 dB: p = 0.010). A main effect of MIA emerged across the increasing intensities in female animals (F(1, 44) = 12.028, p = 0.001, *n*^*p*2^ = 0.215; **Figure 4H**) in which poly (I:C) animals had attenuated PPI compared to saline treated mice. This pattern was apparent at the individual 73, 77, 81, and 85 intensities (73 dB: 0.031; 77 dB: p = 0.002; 81 dB females: p=0.001; 85 dB: p = 0.002; **Figure 4G**).

### 3.4. Effect of MIA and environmental enrichment on hypothalamic gene expression

Given the alterations in pubertal timing we investigated several hypothalamic genes integral to reproductive development and functioning. There was a significant housing effect of the *Gnrh1* gene for male juvenile animals (F(1, 26) =6.220, p = 0.019, *n*^*p* 2^ = 0.193; **Figure 5A**). SD-poly (I:C) males had lower levels than EE-poly (I:C) mice. For female offspring, there was a significant interaction between MIA and housing on this measure (F(1, 28) = 6.471, p = 0.017, *n*^*p* 2^ = 0.188; **Figure 5A**) in that SD-poly (I:C) animals had reduced *Gnrh1* mRNA expression compared to SD-saline (0.047) and EE-poly (I:C) females (p = 0.011) on P32.

**Figure 5.**
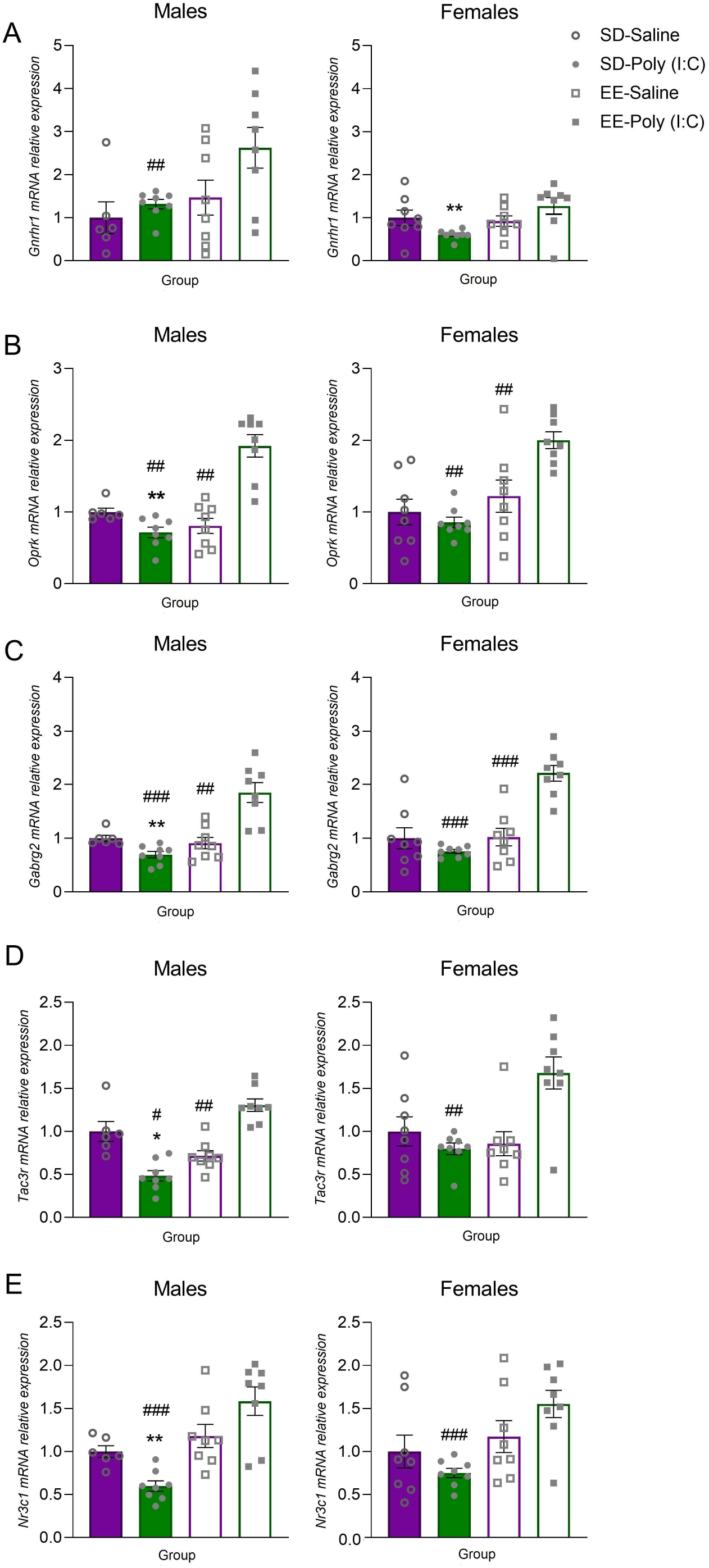
The effects of maternal immune activation (MIA) and environmental enrichment on postnatal day 32 gene expression in the hypothalamus. Male (left) and female (right) offspring levels of **(A)** *Gnrhr1*, **(B)**, *Oprk*, **(C)**, *Gabrg2*, **(D)**, *Tac3r*, **(E)** *Nr3c1*. Data are expressed as mean ± SEM relative to the same-sex SD-control group. **p < 0.01, versus SD-saline; #p < 0.05, ##p < 0.01, ###p <0.001, versus EE-poly (I:C).

Male SD-poly (I:C) animals also had reduced hypothalamic *Oprk* mRNA expression compared to SD-saline (*X*2(1) = 7.350, p = 0.007) and EE-poly (I:C) males (*X*2(1) = 11.294, p = 0.001). Moreover, EE-poly (I:C) male offspring had higher levels of *Oprk* than EE-saline mice in this brain region (*X*2(1) = 10.599, p = 0.001; **Figure 5B**). With respect to female offspring, EE-poly (I:C) mice had higher levels of *Oprk* mRNA than SD-poly ((I:C) *X*2(1) = 6.353, p = 0.012) and EE-saline mice (*X*2(1) = 11.294, p = 0.001; **Figure 5B**).

There was a significant MIA by housing interaction for male *Gabrg2* expression in the male hypothalamus (F(1, 26) = 25.925, p = 0.0001, *n*^*p* 2^ = 0.499; **Figure 5C**). Again, male SD-poly (I:C) offspring had reduced gene expression compared to SD-saline (p = 0.004) and EE-poly (I:C) male mice (p = 0.0001). EE-poly (I:C) males also had higher expression than EE-saline (p = 0.001). There was also a significant MIA by housing interaction for female *Gabrg2* expression in the P32 hypothalamus (F(1, 28) = 23.357, p = 0.0001, *n*^*p*2^ = 0.455). Female EE-poly (I:C) offspring had higher expression than SD-poly (I:C) (p=0.0001) and EE-saline (p=0.0001) female offspring (**Figure 5C**).

Male SD-poly (I:C) offspring had lower hypothalamic *Tac3r* expression compared to both SD-saline (*X*2(1) = 4.817, p = 0.028) and EE-poly (I:C) mice (*X*2(1) = 6.353, p = 0.012). EE-poly (I:C) animals again had higher mRNA expression compared to EE-saline male offspring (*X*2(1) = 4.817, p = 0.028; **Figure 5D**). With respect to female animals, EE-poly (I:C) mice had elevated *Tac3r* expression compared to SD-poly ((I:C) *X*2(1) = 6.893, p = 0.009) and EE-saline mice, but this latter effect was lost to the false discovery rate (*X*2(1) = 4.864, p = 0.027 = N.S; **Figure 5D**).

A MIA by housing interaction for hypothalamic *Nr3c1* (F(1, 26) = 9.452, p = 0.005, *n*^*p* 2^ = 0.267; **Figure 5E**) revealed that SD-poly (I:C) male mice had lower expression of this gene compared to SD-saline (p = 0.003) and EE-poly (I:C) male mice (p = 0.0001). A main effect of housing was found for female mice (F(1, 28) = 9.578, p = 0.004, *n*^*p* 2^ = 0.255; **Figure 5E**) where SD-poly (I:C) mice had lower levels compared to EE-poly (I:C) females (p = 0.0001). With respect to tissue collected on P85, there was a significant interaction between MIA and housing on the level of hypothalamic Kiss1 (F(1, 28) = 6.846, p = 0.014, *n*^*p*2^ = 0.196; **Table 1**) where SD-poly (I:C) males had significantly higher levels of this gene compared to SD-saline (p = 0.023) and EE-poly (I:C) (p = 0.043) males. There were interactions and main effects of sex, MIA, and housing associated with the expression of some additional hypothalamic genes at P32 and P85 which are outlined in **Table 1**.

**Table 1.**
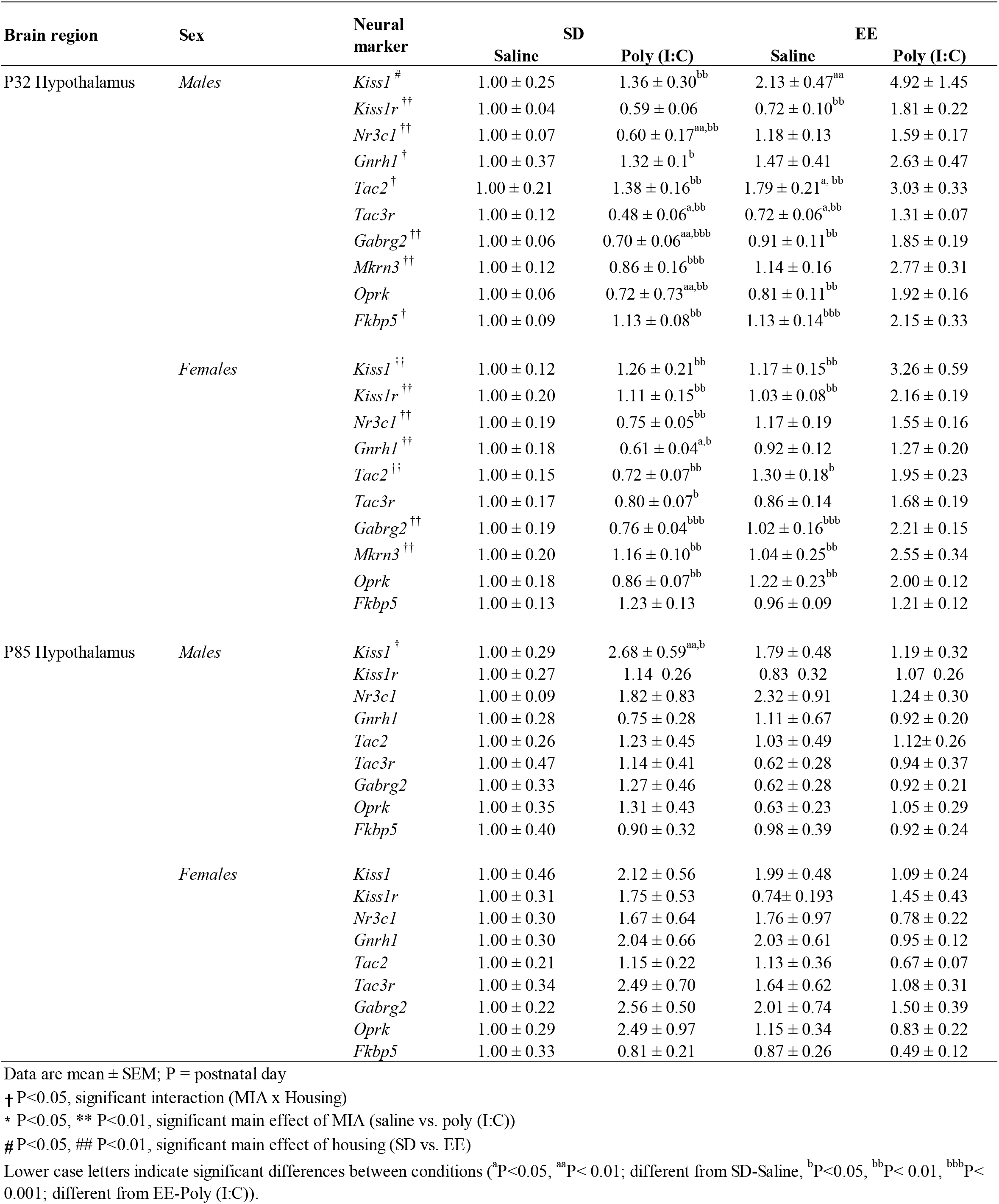
Hypothalamic mRNA expression in offspring exposed to maternal immune activation and housed in SD or EE.

### 3.5. Age of reproductive milestone development is associated with sensory processing

We found consistent associations with age of vaginal opening (r = -0.456, p = 0.001), first estrus (r = -0.0445, p = 0.002), and preputial separation (r = -0.338, p = 0.012) in that the earlier the age the developmental endpoint was reached, the higher the mechanical threshold or sensitivity to the tactile stimulus at P45 (see **Supplementary Data Figure 1**). This relationship reversed in males by P70 where an earlier preputial separation age was associated with a lower mechanical threshold on the Von Frey test (see **Supplementary Data Figure 1**). Additionally, P70 PPI at the 85 dB intensity was negatively correlated with age of preputial separation (r = -0.350, p = 0.025; **See Supplementary Data Figure 1**). There were no significant correlations between the reproductive milestones and P85 gene expression data (see **Supplementary Data Figure 1**).

## 4. Discussion

In the current study, we sought to investigate the impact of MIA on pubertal development, mechanical allodynia, and sensorimotor gating. We also examined the mitigating potential of EE on the effects of MIA. By postnatal day (P) 30, MIA mice in our study displayed altered mechanical allodynia and sensorimotor gating. Both are recognized as symptoms of neurodevelopmental disorders that have been associated with early-life adversity (ELA) and usually manifest following puberty (Le Pen et al., 2006; Salberg et al., 2020) which can be accelerated by early life stress (Gur et al., 2019). Our current study also demonstrated precocious puberty in MIA male and female mouse offspring. This evidence suggests that the accelerated maturation may be a potential mechanism, or catalyst, underlying MIA’s disruptive effects on behavior and neurophysiology.

Precocious puberty following gestational exposure to poly (I:C) is consistent with previous studies showing that MIA and neonatal immune challenges can accelerate the onset of vaginal opening in females (Cakan et al., 2018; Grassi-Oliveira et al., 2016) and that maternal separation can advance preputial separation in males (Yano et al., 2020). This further supports the link between ELA and accelerated pubertal timing. However, there is at least one study showing that ELA (e.g., limited bedding) can delay the onset of vaginal opening in females (Manzano Nieves et al., 2019). As discussed by Manzano Nieves and colleagues (2019), differences in the form of an ELA may partially explain discrepancies between studies. For example, limited access to bedding may not result in a disrupted quality of direct maternal care. In contrast, repeated maternal separations may indicate an increased risk of losing a caregiver, and therefore advance sexual maturation as a strategy to increase reproductive fitness. We previously reported that MIA was associated with poor nest construction quality on P14 (Zhao et al., 2021a) and others have observed MIA to disrupt maternal care quality (Ronovsky et al., 2017). Therefore, immune challenges experienced during gestation, combined with an environment with compromised maternal care, may signal animals to accelerate their puberty timing to maximize reproduction prior to mortality.

Although the molecular mechanisms initiating pubertal timing remain unclear, previous studies suggest that it requires precise coordination of multiple peripheral and central signals that drive the release of downstream hormones from pituitary gonadotrophs, resulting in the development of sex characteristics (Parent et al., 2003). For example, rodent studies showed that puberty is characterized by an increased sensitivity to the stimulatory effects of kisspeptin with respect to GnRH release (Han et al., 2005). Corresponding with pubertal timing is an increase in the number of GnRH neurons expressing *kisspeptin receptor* (Herbison et al., 2010) and *kisspeptin* signaling efficiency (Han et al., 2005). Also, neurokinin B and its receptor, neurokinin-3 receptor (which are encoded by *Tac2* and *Tacr3* mRNAs), play an essential role in normal reproduction (Topaloglu et al., 2009). Moreover, these are upregulated in the brain at puberty (Navarro et al., 2012). However, in the current study, we did not find significant alterations in the expression of most of these genes in the hypothalamus, and surprisingly we found downregulated *Tac3r* and *Gnrh1* respectively in male and female SD-poly (I:C) mice.

Our lab previously demonstrated that early-life adversity, in the form of either litter isolation alongside a neonatal inflammatory challenge or a combination of a Western diet and exposure to limited bedding and nesting protocol, accelerated pubertal timing but decreased the expression of the *GnRH* receptor and *Kiss1* genes respectively (Kentner et al., 2018; Strzelewicz et al., 2019). Our current findings suggest that the effects of some types of early-life stressors on pubertal timing may be mediated through pathways other than direct GnRH-kisspeptin and neurokinin B signaling. For example, in our male SD-poly (I:C) mice, MIA also downregulated *Nr3c1*, a gene that encodes glucocorticoid receptor and plays a critical role in modulating the hypothalamic-pituitary-adrenal (HPA) axis. Downregulated glucocorticoid receptor may alter sensitivity to glucocorticoids, driving excess ACTH secretion and subsequent production of adrenal steroids alongside androgenic and/or mineralocorticoid activity, influencing the timing of puberty onset (Charmandari et al., 2004). It should be noted that for the qPCR assay we evaluated whole hypothalamus, and the critical genes may be expressed in a heterogeneous manner across this brain region. For instance, at the onset of puberty, there is a small population of neurons in the preoptic area that releases GnRH (Goubillon et al., 1999) and the activities of these neurons are regulated by a subpopulation of kisspeptin-neurokinin B-dynorphin neurons in the arcuate nucleus (Lehman et al., 2010). Also, the hypothalamic tissue was collected on P32, and most animals displayed complete vaginal opening and preputial separation before P30. Therefore, we may have missed the critical window for capturing the peak of these essential molecular signals. Our study indicates that it will be important to collect brain tissue specifically timed to pubertal development in future studies. This is particularly relevant for assessing the association between age of achieving reproductive milestones (e.g., vaginal opening, age of first estrus, preputial separation) and gene expression data in brain. In the current study, as these endpoints were evaluated in separate sets of animals, we were not able to relate these variables for a more in-depth analysis. Moreover, a major limitation in this work is the lack of complementary protein or biochemical data to substantiate the biological relevance of our gene expression data. Future studies will need to assess changes in the developmental trajectory of gene expression associated with the reproductive axis, accounting for tissue specificity and changes in protein expression.

The mitigating effects of EE on precocious puberty in female rats have been demonstrated using other early-life stress models (e.g., litter isolation alongside a neonatal inflammatory challenge or limited bedding and nesting; Kentner et al., 2018; Strzelewicz et al., 2019). However, the current study only revealed a modest effect of EE on normalizing the accelerated vaginal opening, and the age of first estrus was even advanced in female EE mice. This discrepancy may be attributed to different types of early-life stressors or the employment of different species between studies. The life-long EE procedure used in the current study consisted of both increased social and physical stimulation, and nesting materials for dams. While the specific contribution of these different EE components is out of the scope of this study, for females housed in larger social groups, one component of the EE procedure, the presence of other females may suppress the occurrence of estrus (Champlin, 1971; Ryan and Schwartz, 1977). This might be one of the factors that also contributes to the modest effects of EE on the timing of vaginal openings. In contrast to the modest effects in females, EE mitigated the advanced day of preputial separation in male mice. The direct comparison between the effects of EE on puberty development in female and male mice has not been reported before, but previous evidence showed sex-differences in the response to the effects of EE at various physiological and behavioral levels (Wood et al., 2010; Zakharova et al., 2012). Our findings indicate that male mice may be more sensitive to EE during pubertal development. At the transcriptional level, EE upregulated the expression of several genes associated with the stress response system and the GnRH-kisspeptin and neurokinin B signaling pathways. Yet, it remains unknown at present the mechanisms for how these signals interplay to mediate EE’s effects on pubertal timing.

We speculate that exposure to EE may alleviate the effects of MIA on accelerated puberty by buffering HPA axis stress responses. This is supported by our findings that EE significantly upregulated hypothalamic *Nr3c1* expression in male and female EE-poly (I:C) exposed mice. At the theoretical level, Boyce and Ellis (2005) proposed a U-shaped, curvilinear relationship between the level of supportiveness versus the level of stress one is exposed to during early life, and how this relates to the development of individual stress reactivity. They propose that individuals who are raised in an environment with intensive, stable caregiving and family support may develop an exaggerated reactivity profile that helps them garner the benefits of such a highly supportive rearing environment. Echoing this notion, our findings of increased *Nr3c1* expression in EE-poly (I:C) challenged animals, combined with observations that MIA animals housed in EE are more responsive to environmental influences on some measures, even compared to non-MIA EE controls (Connors et al., 2014; Núñez Estevez et al., 2020; Zhao et al., 2021a), may reflect their increased susceptibility to benefit from the social and developmental attributes of EE.

Neonatal immune challenge has been associated with the manifestation of hyperalgesia and allodynia in later life (review see Karshikoff et al., 2019)). Combined with similar findings from our current MIA model, this suggests that an immune challenge experienced across different time points in early development can have a long-lasting impact on an animal’s mechanical pain perception. Moreover, the expression of hyperalgesia and allodynia may change across time (Ririe et al., 2003). In one model of post-operative pain, mechanical allodynia thresholds increased across development, with the younger rats (2-4 weeks old) having a significantly lower mechanical threshold than older rats (16 weeks old; (Ririe et al., 2003). In the current study, we also found an age-dependent effect of MIA on von Frey threshold with younger animals being more sensitive than older male and female mice. Interestingly, MIA decreased the allodynia threshold in the SD animals on P30, but increased it on P45, suggesting an age-dependent effect of MIA on pain sensitivity. Therefore, the increased threshold on P45 may reflect a compensatory overshoot for the earlier decreased threshold. Furthermore, the altered allodynia in adolescence was remitted by adulthood, which is consistent to previous finding showing that male rat offspring exposed to valproic acid prenatally displayed lower mechanical allodynia as adolescents, which was recovered when the animals reached adulthood (Schneider et al., 2006). Indeed, our correlational analyses showed that male and female mice that demonstrated an earlier age of vaginal opening, first estrus, and/or preputial separation, had higher mechanical allodynia thresholds on P45. This association was reversed in male animals by P70 and absent in females. The specific mechanisms underlying this temporal expression is still unclear but the high correlation on this outcome suggests that accelerated puberty may be a potential driver for changes in tactile sensitivity, at least at defined windows of development, e.g., adolescence. Our own work has previously demonstrated that in adolescence, rats that were exposed neonatally to lipopolysaccharide displayed disrupted social behavior which were remitted in adulthood (MacRae et al., 2015). Our findings suggest that adolescence may be an important developmental stage during which the effects of MIA on allodynia emerge, even if only transiently.

Consistent to a previous finding that EE rescues increased pain sensitivity in young rats prenatally exposed to valproic acid (Schneider et al., 2006), the current study also revealed a protective effect of EE on the disrupted allodynia following MIA. However, we were surprised to observe that on P30, EE-saline females also had a lower allodynia threshold than SD-saline controls. This seems contrary to the general beneficial effects of EE. One possible explanation is sex differences in the response to EE, which has not been well studied in previous EE research which has tended to focus on males (Kentner et al., 2019b; Kentner et al., 2021). Of the work that has explored sex-differences, some studies in rats found an overall greater effects of EE for males than for females, in terms of locomotor activity (Elliott and Grunberg, 2005), social exploration (Peña et al., 2006), and social memory. The latter was assessed by social discrimination task which was improved by EE in males but impaired in females (Peña et al., 2009). At the transcriptional level, our results showed that poly (I:C) downregulated, while EE rescued, the mRNA expression of opioid receptor kappa 1, an important component of the endogenous opioid system which plays an essential role in antinociception (Ossipov et al., 2010). This finding suggests that MIA’s disruptive effects (and EE’s resulting protective effect) on mechanical allodynia may be mediated, at least partially, through modulation of the opioid system. Reduced PPI values usually signify deficits in sensorimotor gating, which are correlated with several positive and negative symptoms of schizophrenia (Braff et al., 1999; San-Martin et al., 2020). Similar to the manifestation of other symptoms of this neurodevelopmental disorder, deficits in sensorimotor gating usually occur after puberty (Le Pen et al., 2006). In the current study, we did not observe an overall disruptive effect of MIA on PPI; instead, very specific impairments of PPI appeared to depend on age, sex and the particular prepulse intensities employed. With respect to the MIA literature, despite reports that poly (I:C) treatment results in a general disruptive effect on PPI in offspring (Howland et al., 2012; Meehan et al., 2017; Meyer et al., 2010; Shi et al., 2003; Wolff and Bilkey, 2008), its efficacy appears to be dependent on various factors and the findings are not consistent. For example, Wolff and Bilkey (2010) found that regardless of the presence of weight loss in MIA dams following injection, the offspring showed impaired PPI in both the juvenile period and adulthood. However, others have observed that only the offspring of dams that lost weight following poly (I:C) injection showed altered PPI responses (Vorhees et al., 2015). Moreover, other reports suggest that poly (I:C) administration affects PPI only in adulthood, but not in juveniles (Ding et al., 2019; Ozawa et al., 2006). Furthermore, different molecular weights and dosages of poly (I:C) can give rise to varied efficacy on different aspects of behavioral/physiological outputs (Careaga et al., 2018; Kowash et al., 2019; Mueller et al., 2019). All this evidence adds to the complexity of MIA’s effects on sensorimotor gating ability and highlights the importance of optimizing MIA protocols for studying sensorimotor gating deficits.

In the current study, we used 20 mg/kg of low molecular weight poly (I:C) and this dosage of administration implemented at a mid-gestational stage only resulted in a modest attenuation of PPI compared to saline treated mice across the increasing prepulse intensities. Furthermore, we found exposure to life-long EE exerted a sex-and age-dependent enhancing effect on the PPI, which is reflected by our results showing the increased PPI in the EE group compared to the SD group on P30 and P70 in only males. Previous studies have reported inconsistent impacts of EE exposure on sensorimotor gating. For example, pre-and postweaning EE housing mitigated the diminished PPI in adolescent male rats prenatally treated with valproic acid (Schneider et al., 2006) while EE exposure starting at weaning and continuing into adulthood was associated with decreased or no effect on PPI (Hendershott et al., 2016; Peña et al., 2009). Factors such as the duration and the starting age of EE exposure may explain these discrepancies.

To summarize, the present study revealed disruptive effects of MIA on pubertal development, mechanical allodynia, sensorimotor gating, and hypothalalmic mRNA expression related to sexual maturation and HPA functioning. These findings suggest that accelerated puberty is associated with exposure to early life adversity and may contribute to a wide range of MIA-related physical and mental health complications. Furthermore, we found a general beneficial effect of EE on the abovementioned disruptions induced by MIA, suggesting potential translational utility of EE in mitigating negative consequences of early life experiences. Compared to pharmacotherapies, EE is a non-invasive intervention and may serve as a prevention strategy for pregnant women who experience severe inflammatory insults. Our findings provide further evidence for the variable effects that MIA has in the etiology of neurodevelopmental disorders and the potential therapeutic or preventive role of the environment in mitigating some of the consequences of early life adversity.

## Funding and Disclosure

This work was supported by the National Institute of Mental Health (R15MH114035 to ACK). The authors wish to thank Ryland Roderick and Madeline Puracchio for technical support during the early phases of this study. The authors would also like to thank the MCPHS University School of Pharmacy and School of Arts & Sciences for their continual support. The content is solely the responsibility of the authors and does not necessarily represent the official views of any of the financial supporters.

## Author Contributions

X.Z., R.M., & M.E., ran the experiments; X.Z., & A.C.K. analyzed and interpreted the data, and wrote the manuscript; A.C.K., designed and supervised the study and made the figures.

## Declaration of Competing Interest

The authors declare that they have no known competing financial interests or personal relationships that could have appeared to influence the work reported in this paper.

## Data Availability Statement

All data are available upon request to the authors.

**Supplementary Figure 1. Correlational data between puberty milestones, mechanical allodynia, and PPI**. Association between P45 mechanical allodynia and female age A) of vaginal opening, B) at first estrus, and male age C) at preputial separation. Associations between age of male preputial separation and P70 D) mechanical allodynia and E) P70 PPI at an 85 dB intensity.

**Supplementary Table S1.** Maternal Immune Activation Model Reporting Guidelines Checklist.

## Notes

### Competing Interest Statement

The authors have declared no competing interest.

