## Supplementary figures and images for "Maternal immune activation accelerates puberty initiation and alters mechanical allodynia in male and female C57BL6/J mice"

### Supplementary Data Figure 1

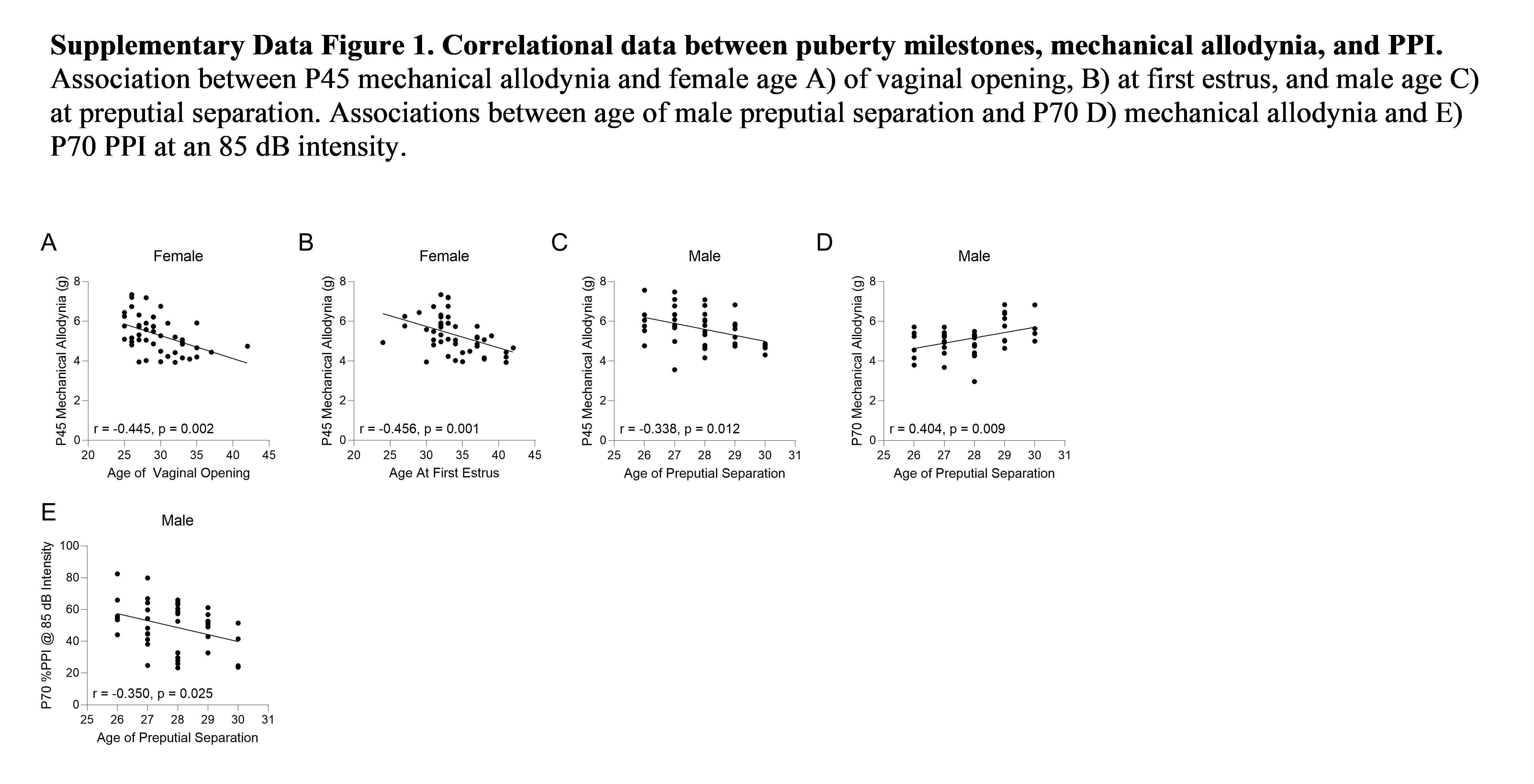
